# MeDUsA: A novel system for automated axon quantification to evaluate neuroaxonal degeneration

**DOI:** 10.1101/2021.10.25.465674

**Authors:** Yohei Nitta, Hiroki Kawai, Jiro Osaka, Satoko Hakeda-Suzuki, Yoshitaka Nagai, Karolína Doubková, Takashi Suzuki, Gaia Tavosanis, Atsushi Sugie

## Abstract

**Background:** *Drosophila* is an excellent model organism for studying human neurodegenerative diseases (NDs), and the rough eye phenotype (REP) assay is a convenient experimental system for analysing the toxicity of ectopically expressed human disease genes. However, the association between REP and axonal degeneration, an early sign of ND, remains unclear. To address this question, we developed a method to evaluate axonal degeneration by quantifying the number of retinal R7 axons in *Drosophila*; however, it requires expertise and is time-consuming. Therefore, there is a need for an easy-to-use software that can automatically quantify the axonal degeneration.

**Result:** We created MeDUsA (a ‘<u>me</u>thod for the quantification of <u>d</u>egeneration <u>us</u>ing fly <u>a</u>xons’), which is a standalone executable computer program based on Python that combines a pre-trained deep-learning masking tool with an axon terminal counting tool. This software automatically quantifies the number of axons from a confocal z-stack image series. Using this software, we have demonstrated for the first time directly that axons degenerate when the causative factors of NDs (αSyn, Tau, TDP-43, HTT) were expressed in the *Drosophila* eye. Furthermore, we compared axonal toxicity of the representative causative genes of NDs and their pathological alleles with REP and found no significant correlation between them.

**Conclusions:** MeDUsA rapidly and accurately quantifies axons in *Drosophila* eye. By simplifying and automating time-consuming manual efforts requiring significant expertise, it enables large-scale, complex research efforts on axonal degeneration, such as screening to identify genes or drugs that mediate axonal toxicity caused by ND disease proteins.

## Background

Neurodegenerative diseases (NDs) are disorders in which certain groups of neurons in the brain and spinal cord involved in cognitive and motor function are gradually lost. Molecular genetic studies have identified causative genes and risk factors and elucidated the mechanisms of pathogenesis at the molecular level. These findings revealed that structural defects and aggregation of disease-associated proteins underlie neurodegenerative processes [1]. It has become available to examine the effects of novel mutations found in human diseases like NDs, whether they result in the loss of gene function or gain of toxic function, using various model organisms. Among them, *Drosophila* has various advantages as a neuronal disease model. For example, 1) gene function can be analysed without strongly considering the compensation of gene function by duplication since there are relatively few duplicated genes in the genome; 2) research sample sizes can be large because individuals are small, inexpensive and easy to breed; 3) the short life cycle allows rapid genetic analysis and 4) the organism has a compact brain, making it possible to analyse at the level of neural circuitry and behaviour necessary for higher functions such as learning, memory and sleep. Taking advantage of these features, *Drosophila* is widely used in studies on human diseases. Further, it has been reported that the expression of a human disease-associated protein in *Drosophila* induces toxicity even in flies [2–6], demonstrating the conservation of molecular mechanisms between humans and flies.

Trinucleotide repeat disorders are human diseases caused by the expansion of CAG repeats in the protein-coding regions of causative genes. Spinocerebellar ataxia type 3 (SCA3), also known as Machado-Joseph disease, is a neurodegenerative disease caused by repeated elongation of glutamine. In *Drosophila*, the expression of these extended polyglutamine repeats not only formed inclusion bodies similar to those in humans, but also caused degeneration [2]. CAG repeats are also found in Huntingtin (HTT), the gene responsible for Huntington’s disease, which is another autosomal dominant neurodegenerative disease. In experiments in which polyglutamine-extended HTT was expressed in the photoreceptors of *Drosophila*, inclusion bodies formed and the polyglutamine-extended HTT induced neurodegeneration [3]. Additionally, in Parkinson’s disease (PD), a neurodegenerative disorder characterised by the loss of dopaminergic neurons in the substantia nigra, formation of Lewy bodies and impaired motility, expression of the causative gene synuclein alpha (SNCA) in *Drosophila* caused the loss of dopaminergic neurons, the formation of fibrous intraneuronal inclusions containing Alpha-synuclein (αSyn) and motor dysfunction [4]. Furthermore, the expression of the microtubule-associated protein Tau, which is involved in Alzheimer’s disease, in all neurons of *Drosophila* led to the observation of progressive neurodegeneration [5]. Thus, *Drosophila* models expressing human disease-causing genes can reproduce the characteristics of human diseases, thereby enabling powerful genetic approaches to study various NDs such as polyglutamine disease, synucleinopathy and tauopathy. Using these ND models, large genetic screens can be performed to explore unknown protein networks in which disease-causing proteins interact. The homologues of the candidate network members can then be identified in the human genome to determine whether they are susceptibility genes of the disease of interest. In one example of a disease study of amyotrophic lateral sclerosis (ALS), genetic screening using this experimental paradigm identified that ATXN2 is involved in ALS pathogenesis [7]. Further, suppression of the ATXN2 *Drosophila* homologue reduced the toxicity of TDP-43, which is a DNA/RNA-binding protein implicated in several NDs. The discovery of ATXN2 accumulation in the spinal cord of human ALS is but one example revealed using this screening approach in *Drosophila*.

The rough eye phenotype (REP), which is frequently used to investigate genetic interactions in *Drosophila*, is also used in disease research as a convenient and quick way to assess the toxicity of ectopically expressed genes. The phenotypic assay uses the Gal4/UAS method [8] to evaluate the toxicity of a gene of interest by expressing it specifically in the eye using the eye-specific *Gal4*, *GMR-Gal4* and observing eye structure. In fact, this fly eye assay has identified several modifiers that inhibit or enhance the toxicity caused by pathogenic factors such as Tau [9–11], αSyn [12–15], TDP-43 [16, 17] and polyglutamated HTT [18–20]. Moreover, drug screenings can be performed by adding compounds to fly food, and compounds that reduce disease toxicity can be identified [21, 22]. Although REP has been evaluated qualitatively in most previous studies, several research groups have recently reported methods for the quantitative evaluation of REP [23, 24]. However, it remains unclear whether REP reflects neurodegeneration completely as the formation and geometric defects of cell clusters are assessed only within the ommatidium, which contains cone and pigment cells. To assess neurodegeneration more fully in *Drosophila*, several other parameters related to neurodegeneration, as measured by protein aggregation number, vacuolar size and number and retinal thickness, have been previously developed [25]. However, these systems also indirectly observe neurodegeneration, and therefore, a direct quantitative method for accurately measuring neurodegeneration is required.

Axonal degeneration is a representative pathology of neurodegeneration, and the magnitude of neurodegeneration can be accurately evaluated by its quantification. To date, there are few methods for quantifying axonal degeneration [26, 27]. Further, they are time-consuming, requiring the manual quantification of axonal number or depend on the expertise of experimenters to accurately classify the degree of degeneration. However, biological images contain noise, and variable signal intensities are often observed among samples. Additionally, the three-dimensional (3D) structure and angle are never uniform between images. Thus, to recognise the semantic region from diverse image data, the eye of a trained researcher can flexibly and accurately extract specific phenomena to be analysed, but such time-consuming methods are not suitable for screening to identify genes or compounds that modify the pathology and requires quantifying large numbers of samples. Therefore, robust image processing systems other than the human eye are required to evaluate axonal degeneration simply and quickly. In recent years, image processing technology has made remarkable progress, especially the development of deep-learning technology using convolutional neural networks (CNNs), which has greatly advanced the field of image recognition, and is also true in the field of biological images [28]. In addition, although conventional image processing can extract signal regions, it is difficult to extract semantic regions, but CNN has demonstrated high performance in semantic domain segmentation. In particular, U-Net [29] is an architecture of CNNs designed for biological image analysis, and U-Net and its derived architectures have been used for segmentation tasks in the area of biological image analysis with great success [30–33].

In an accompanying study, we developed a novel method to directly quantify axonal degeneration using R7, a photoreceptor neuron type in *Drosophila*, as a model [34]. In this method, axon terminals are manually excised from the confocal microscope z-stack image series, and degeneration is quantified by manual counting the number of axon terminals. This allowed quantifying even minor axon losses; therefore, even very early stages of axonal degeneration phenomena. Nonetheless, its throughput was not sufficient for larger scale screens. To automate the method, we established here a novel software package called ‘<u>me</u>thod for the quantification of <u>d</u>egeneration <u>us</u>ing fly <u>a</u>xons’ (MeDUsA) by combining deep learning with a Python-based counting system. Using this software, we assessed the effects on axons among the causative genes of several NDs and found that they exhibited axonal degeneration. Additionally, no significant correlation was detected between the number of axons and the REP. MeDUsA provides direct and rapid quantitation of axonal degeneration in *Drosophila*, making it a powerful tool that can be used in disease and developmental studies.

## Results

### Rough eye phenotype is insufficient to speculate on gene effects in axonal degeneration

The Rough eye phenotype (REP) assay has been extensively used to study neurodegenerations (NDs) [35]; however, there is uncertainty whether REP accurately reflects axonal degeneration. We performed a modifier genetic screening designed to identify genes that modulate the toxicity of TDP-43 by observing retinal and axonal degeneration phenotypes. The *Drosophila* photoreceptors R7 and R8 project their axons directly from the compound eye retina through the primary optic ganglion lamina to the secondary optic ganglion medulla (Fig. 1A). For this screening, a fly line with eye-specific expression of TDP-43^G298S^, which is an ALS-associated mutation of TDP-43 [36], using *GMR-Gal4* [37, 38] was generated. These flies were crossed with 99 candidate RNA interference (RNAi) lines (Fig. 1B; see Materials and Methods). In the first screening round, we observed the eye phenotype because the eye-specific expression of TDP-43^G298S^ causes REP as previously reported [39]. As a result, 14 of 99 RNAi lines were identified that suppress REPs (Fig. 1B, 1C). Next, we evaluated the axonal degeneration of R8 retinal axons, as the second screening round (Fig. 1B, 1C). To quantify the axonal degeneration, we calculated the ratio of degenerated axons in the part of the optic lobe that is easy to observe each axon. We classified an axon as degenerated when the axon was fragmented. Eye-specific expression of TDP-43 displayed axonal degeneration of R8 axons one day after eclosion (Fig. 1C). We expected that REPs identified in the first screening round would be consistent with the morphology of the R8 axons under the assumption that RNAi lines which suppressed REPs rescued the axonal degeneration. Surprisingly, the REP results did not always match those of axonal degeneration. Knockdown of *Dsk*, a neuropeptide-encoding gene identified in only crustaceans and insects, in the background of TDP-43 expression strongly suppressed REP; however, *Dsk* knockdown significantly promoted axonal degeneration (Fig. 1C; quantified in 1D). In addition, the knockdown of 10 genes did not affect axonal degeneration, whereas 3 genes (*mle*, *faf* and *caz*) rescued the degeneration (Fig. 1C; quantified in 1D). Thus, although the REP assay is a simple and powerful method for assessing toxicity, it is insufficient to evaluate axonal degeneration. Further, the quantitative method we used to assess degeneration in this screening was also insufficient for precise quantification because it was a subjective determination whether an axon was degenerated (i.e. fragmented) or intact. Other limitations include the fact that axons which are completely lost cannot be counted and all R8 axons were not counted from the dorsal view due to limits of the confocal microscope scanning time and depth at which the sample could be viewed cleanly. Additionally, depending on the person performing the method and region being quantified, the results would display a high degree of variability. Therefore, we developed an automatic system, MeDUsA for the unbiased quantitative evaluation of axonal degeneration in *Drosophila*.

**Figure 1.**
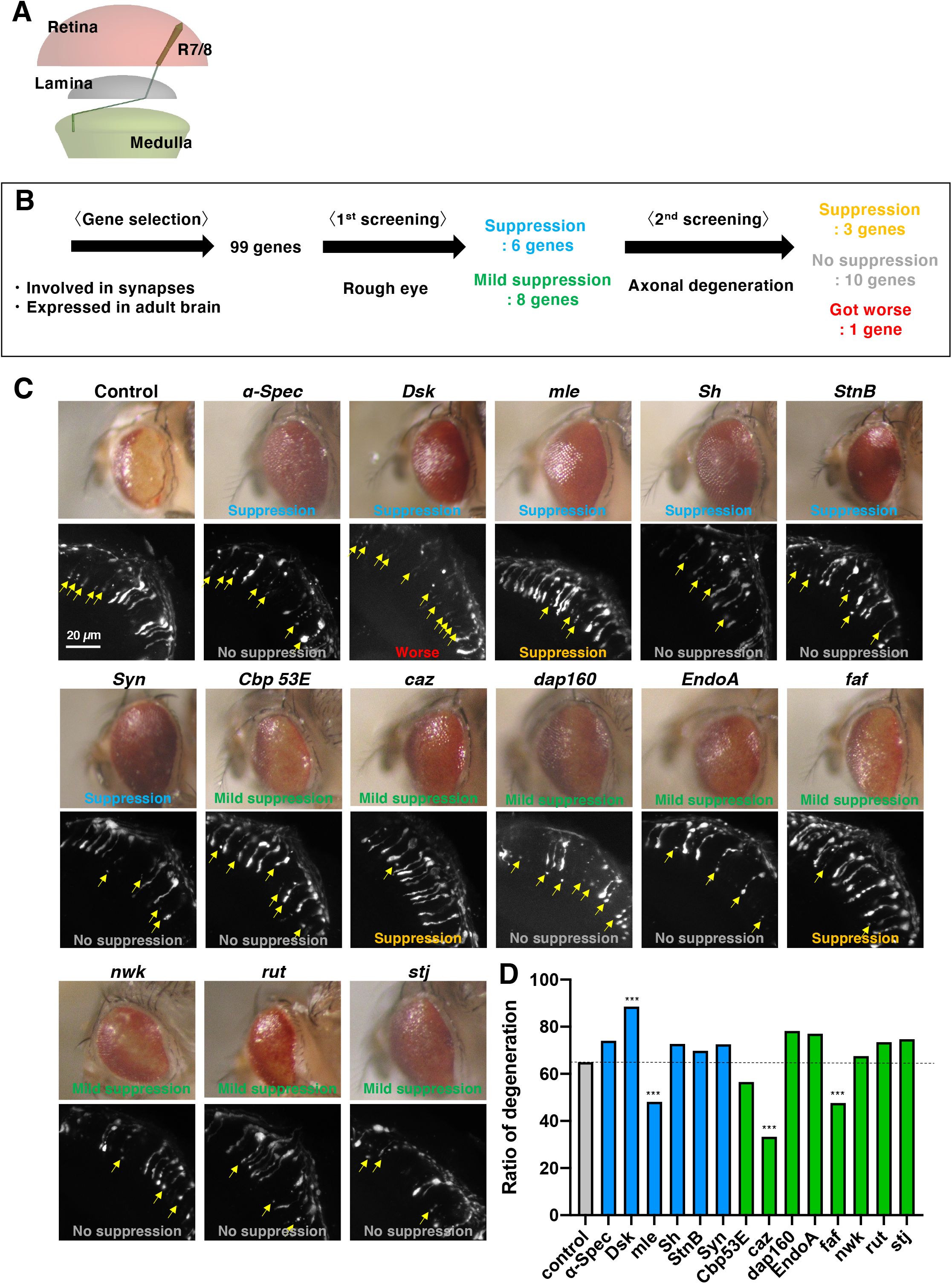
Rough eye and axonal degeneration phenotypes for evaluating neurodegeneration. (A) Dorsal schematic of the visual system in *Drosophila*. The axons of photoreceptors R7 and R8 project from the retina through the lamina to the medulla. (B) The process of exploring factors that mitigate TDP-43 toxicity using a combination of rough eye and axonal degeneration phenotypic observations. (C) Candidate genes that suppress the rough eye phenotype (REP) and their involvement in axonal toxicity. In knockdown screening, six genes suppressed REP and eight genes mildly suppressed it. However, the degree of REP and severity of R axon degeneration were not consistent. Yellow arrows indicate fragmented axons. Scale bar = 20 *μm*. (D) Quantification of the ratio of axonal degeneration. \*\*\**p < 0.001*. Chi-square test was performed between the control and each knockdown.

### The process flow for using MeDUsA

To quantify R7 axons using MeDUsA, the first step is to prepare samples of the *Drosophila* brain (Fig. 2A and 2B). Dissection and immunostaining were performed as previously described [34, 40].

**Figure 2.**
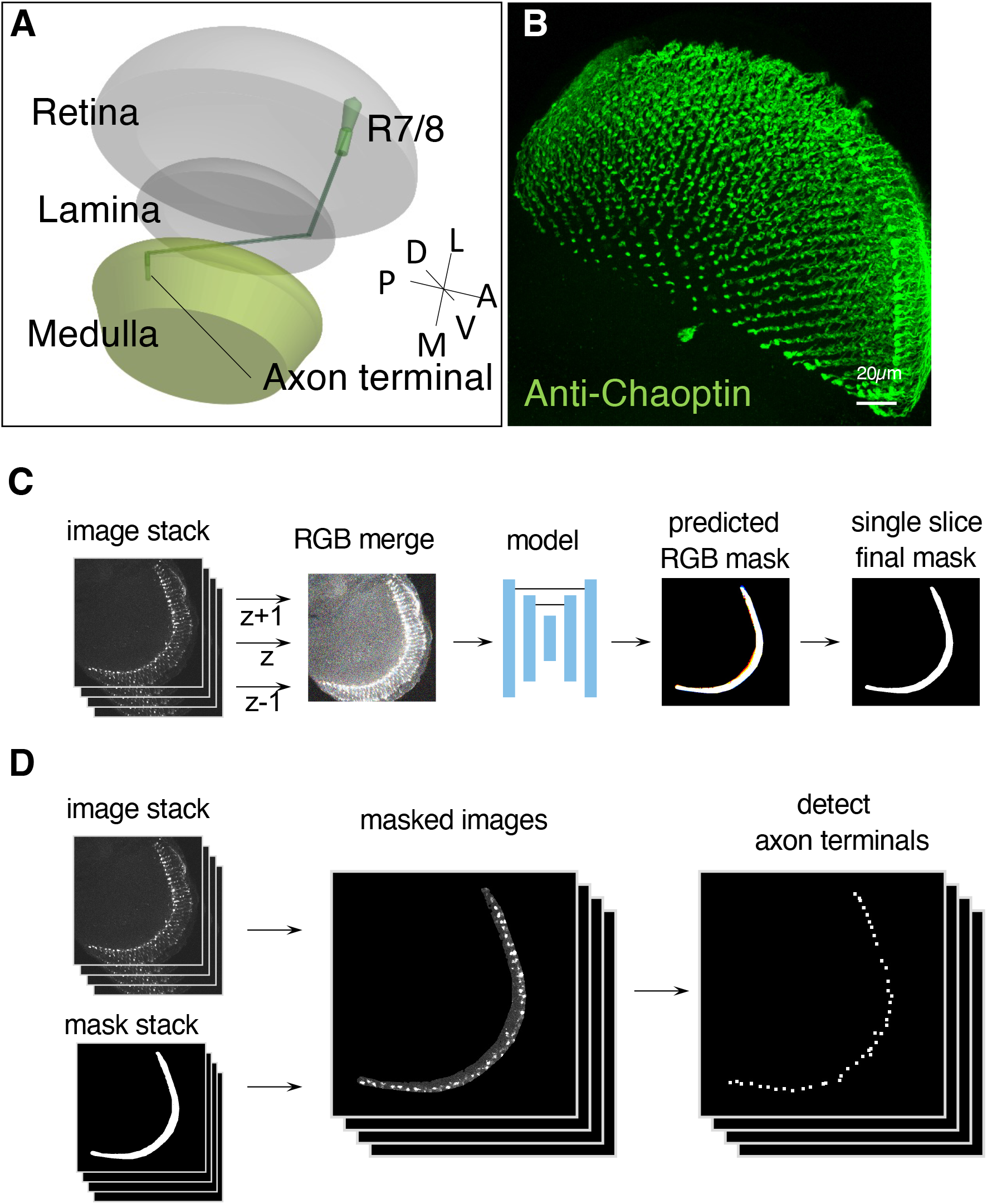
Sample Preparation and Processing flow of MeDUsA. (A) Schematic of the visual system in *Drosophila*. (B) All R7 and R8 axon terminals projecting to the medulla. R axons were stained with anti-Chaoptin, which is a photoreceptor-specific antibody (green). Scale bar = 20 *μm*. (C) Process flow of generating a surface mask. First, for each slice of the z-stack image, three slices (z − 1, z and z + 1) are merged by adding the previous and next slice to create an RGB image. The RGB image is inputted into the training model to generate the RGB mask. Finally, only the channel corresponding to the central slice is extracted to obtain the final mask. (D) Process for axon terminal detection. Using the obtained surface mask stack and original image stack, the surface mask region is extracted from the original image for each slice, and only the signal at the axon end is extracted. Axon terminals are then detected and quantified from the obtained 3D volume.

In the second step, axon termini were identified and quantified. For this purpose, we developed MeDUsA to enable non-experts to accurately evaluate axonal degeneration and to save time. MeDUsA utilises a combination of tools available in Python that allows the task of masking axons to be performed by pre-trained deep learning, followed by automatic counting of axon terminals after masking. This software enables researchers to quantify the number of R7 axons readily and quickly, taking 50 seconds per sample on a general desktop workstation (CPU: Intel Core i7 9800X 3.8 GHz, RAM: DDR4 128 GB).

To create a model that generates a mask of the surface area, we trained a 2D-U-Net architecture (Fig. 2C). However, to determine the surface area, it is necessary to infer it from the axon terminal signals in the sparsity, and there are areas where it is difficult to determine the surface area with a single z-slice. Therefore, we trained the 2D-U-Net to generate a surface mask of three slices by inputting three slices as three channels, including the slice before and after the slice of interest. Then, during inference, only the mask of the centre slice was used. The training and testing datasets included both normal and abnormal images. A total of 16,114 images in 199 samples were used for training and 1,375 images in 16 samples were used for testing. By using three channels, the dice score improved from 0.815 to 0.847 compared to using a single channel.

After mask generation and axon terminal extraction, the number of axonal terminals was counted automatically (Fig. 2D). To do this, we first filtered the regional maxima to remove background, followed by binarisation using adaptive thresholding. The surface area was then extracted using mask images. We performed a Euclidean distance transformation to obtain peaks, which were used as seeds to perform 3D watershed to obtain each axon terminal candidate. Finally, candidates below 20 voxels (equivalent to a radius of about 1.68 *μm*) were eliminated, and the remaining ones were counted as axon terminals. This process was fully automated and allowed us to stably detect axon terminals without adjusting parameters for each sample.

### Ectopic expression of causative genes of NDs causes axonal degeneration in R7 neurons

Using our MeDUsA, we evaluated the effect of mutations in proteins responsible for NDs on axonal degeneration in *Drosophila*. We expressed either wild-type or well-known mutant alleles of human causative genes for neurodegenerative diseases (αSyn, Tau, TDP-43 and HTT) in photoreceptor axons using *GMR-Gal4* and predicted the number of axonal terminals in 1-day-old adults. The ectopic expression of wild-type αSyn, which is a causative gene of PD [41], in photoreceptor axons did not show a significant reduction in axonal number compared to control, whereas the expression of A53T-mutated αSyn, a well-known pathogenic allele associated with familial PD [42], caused a significant reduction in the number of axons compared to control (Fig. 3A–C; quantified in 3K). However, there was no significance between wild-type and A53T-mutated αSyn. Next, we found that expression of wild-type Tau (Tau^WT^), which is implicated in Alzheimer’s disease, caused significant axonal degeneration, and the expression of Tau with the R406W mutation, which is a missense mutation identified in families diagnosed with frontotemporal dementia and parkinsonism linked to chromosome 17 [43], enhanced the degeneration (Fig. 3D, 3E; quantified in 3K). Interestingly, the expression of Tau with the S2A mutation, a mutation with impaired phosphorylation capabilities, significantly suppressed axonal degeneration compared to Tau^WT^ (Fig. 3F), indicating that phosphorylated Tau exhibits toxicity. Next, we evaluated the expression of both wild-type and A315T-mutated [44] TDP-43, which has been identified as the major disease protein in ALS, in photoreceptor axons, and found that both displayed axonal degeneration, although degeneration was milder in the A315T mutant than in wild-type TDP-43 (Fig. 3G, 3H; quantified in 3K). Finally, we found that ectopic expression of wild-type HTT (HTT^Q0^), the causative gene of Huntington’s disease, did not cause axonal degeneration, whereas the expression of HTT with a pathogenic polyQ tract of 128 repeats (HTT^Q128^) significantly reduced the number of R7 axons (Fig. 3I, 3J; quantified in 3K). These results show that the ectopic expression of causative genes of human NDs induced axonal degeneration in *Drosophila*, and well-known pathogenic mutations enhanced degeneration, except for TDP-43. Our findings demonstrate that the toxicity of various human disease causative proteins can be reliably assessed across species using our quantitative axonal degeneration fly model.

**Figure 3.**
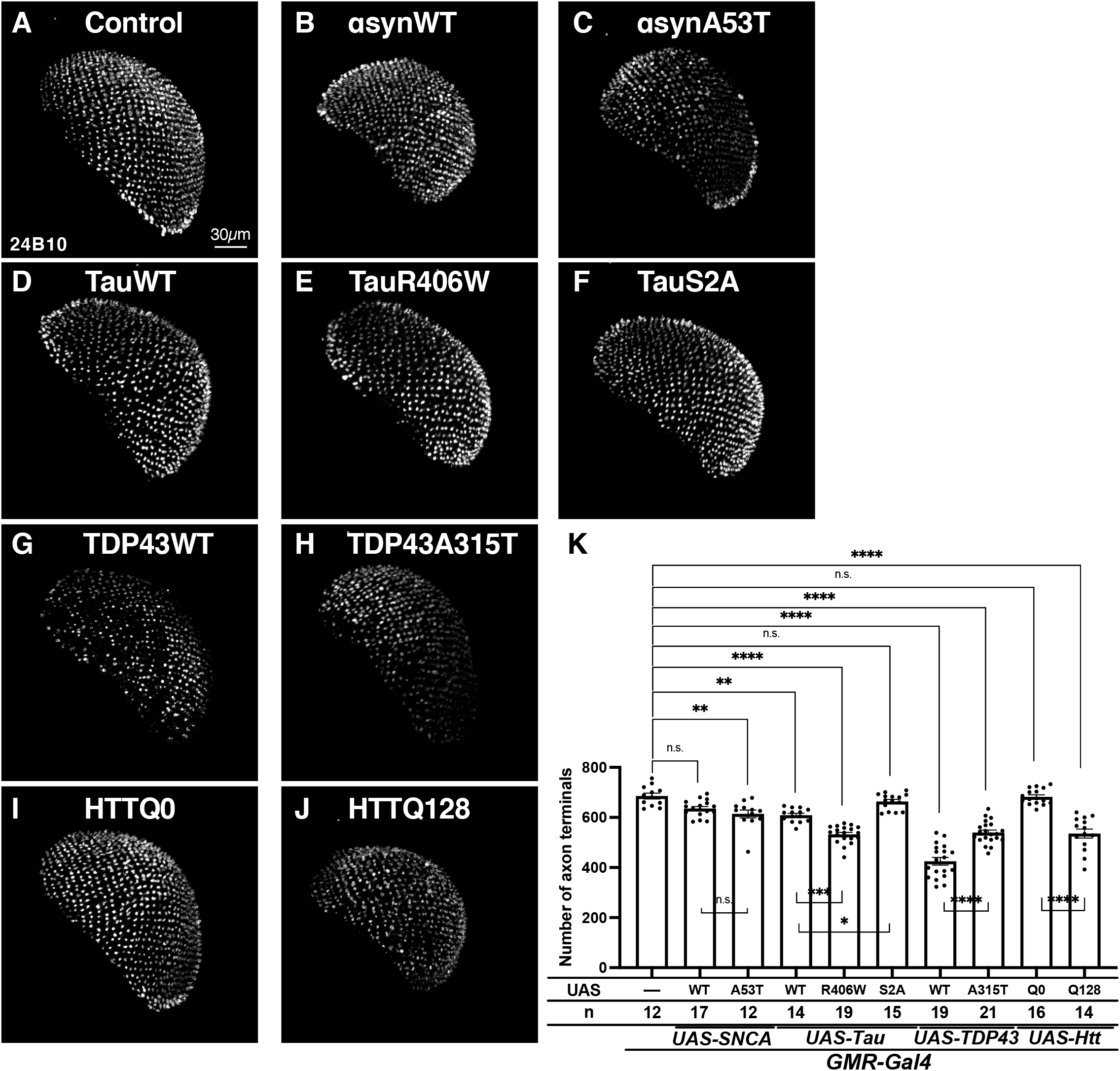
Toxicity evaluation of representative ND causative factors using MeDUsA. (A–J) The R axon terminals stained with anti-Chaoptin and extracted using MeDUsA. (A) Control and photoreceptor expression of (B) αSyn^WT^, (C) αSyn^A53T^, (D) Tau^WT^, (E) Tau^R406W^, (F) Tau^S2A^, (G) TDP-43^WT^, (H) TDP-43^A315T^, (I) HTT^Q0^ and (J) HTT^Q128^. Scale bar = 20 μm. (K) The number of axons expressing each pathogenic factor was predicted. \*\*\*\**p* < 0.0001, \*\*\**p* < 0.001 and ns (*p* > 0.05). Data were analysed using multiple comparison ANOVA with Tukey–Kramer *post hoc* tests. Error bars show the standard error of the mean.

To evaluate the performance of MeDUsA, we quantified the axonal number of the same sample set manually (Fig. 4A). This manual method enabled us to carry out precise quantification of axonal number; however, manual quantification is time-consuming and several parameters have to be adjusted for each sample to detect axon terminals. The MeDUsA measurements were lower than the manual measurements (Fig. 4B). This is due to the severity of the setting (Fig. 2D) for recognising the axon terminal in automated quantification. If the setting is further set loose, the false positive count increases. At present, this is the limitation of the MeDUsA. Nevertheless, the system showed a quadratic weighted kappa score (κ2) of 0.724 and a significantly strong positive correlation with manual counting (individual values, *R^2^* = 0.887, *p* < 0.001, Fig. 4C; mean value per genotype, *R^2^* = 0.944, *p* < 0.001, Fig. 4D). Taken together, these findings demonstrate that MeDUsA automatically and rapidly counts axonal number in a preparation accessible for genetic screening.

**Figure 4.**
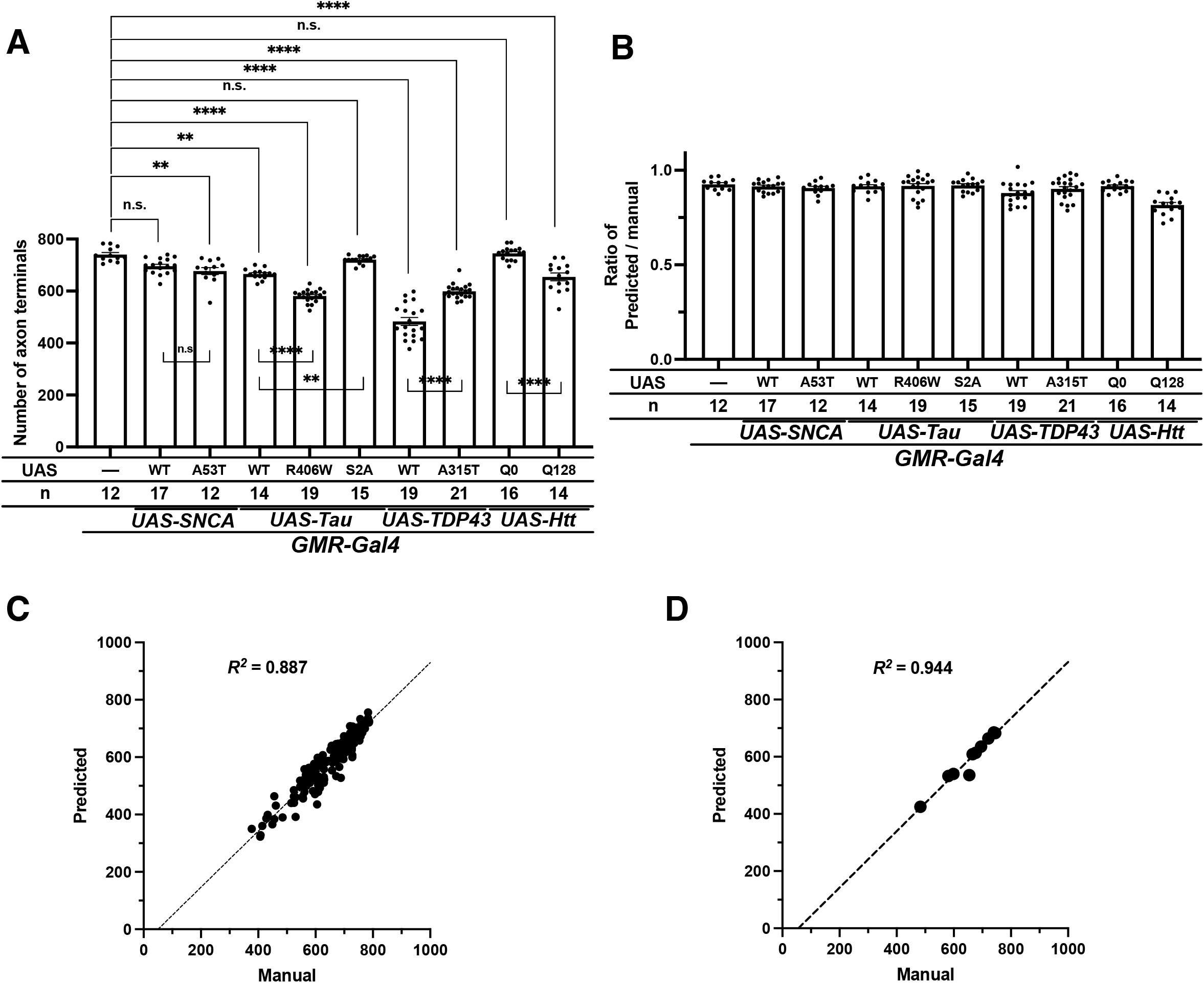
Comparison between manual quantification and the MeDUsA system. (A) The axonal numbers quantified by manual measurements using the same dataset shown in Figure 3K. \*\*\*\**p* < 0.00001, \*\*\**p* < 0.001, \*\**p* < 0.01, \**p* < 0.05 and ns (*p* > 0.05). Data were analysed using multiple comparison ANOVA with Tukey–Kramer post hoc tests. Error bars show the standard error of the mean. (B) The ratio of the axonal number predicted by MeDUsA to those measured manually for each genotype. (C) The correlation between the individual values measured manually and the individual values predicted by MeDUsA. (D) The correlation between the average of each genotype measured manually versus those predicted by MeDUsA. R^2^, coefficient of determination.

### REP does not match the axonal degeneration phenotype

Next, we evaluated the correlation between axonal degeneration phenotype and the REP. To quantify REP severity, we used Flynotyper that calculates the phenotypic score from the disarray of the ommatidia [24]. We expressed the same set of causative genes for NDs in the eye using *GMR-Gal4* as in axonal degeneration experiments and quantified the degree of phenotypic severity of eye phenotypes in 1-day-old adults. We found that the expression of either the wild-type or pathogenic allele of αSyn did not significantly increase the phenotypic score compared to control (Fig. 5A–C; quantified in 5K), although the expression of the pathogenic alleles of αSyn caused axonal degeneration (Fig. 3K). In contrast, REP severity was consistent with the reduction of axonal number when the wild-type and mutant alleles of Tau were expressed (Fig. 5D–F; quantified in 5K). Wild-type and R406W-mutated Tau, both of which caused axonal degeneration, showed a significantly increased phenotypic score, whereas the S2A mutation did not display any reduction in axonal number nor corresponding score increase. Similar to αSyn, we found that TDP-43 expression showed an obvious discrepancy between the two phenotypes. As described earlier, wild-type TDP-43 is more toxic than A315T-mutated TDP-43 on axonal degeneration (Fig. 3K). However, the A315T mutation caused a significant increase in the REP score compared with wild-type TDP-43 (Fig. 5G, 5H; quantified in 5K). HTT expression also exhibited differences between axon and eye phenotypes. In contrast to axonal degeneration phenotypes, the ectopic expression of HTT^Q128^ did not cause REPs (Fig. 5I, 5J; quantified in 5K). Finally, we determined the correlation between the average axonal number and average phenotypic score of rough eye in each genotype, and found a negative correlation, but the correlation was not statistically significant (*R* = −0.467, *p* = 0.174). Taken together, our findings show that REP scoring is not exactly consistent with the degree of axonal degeneration.

**Figure 5.**
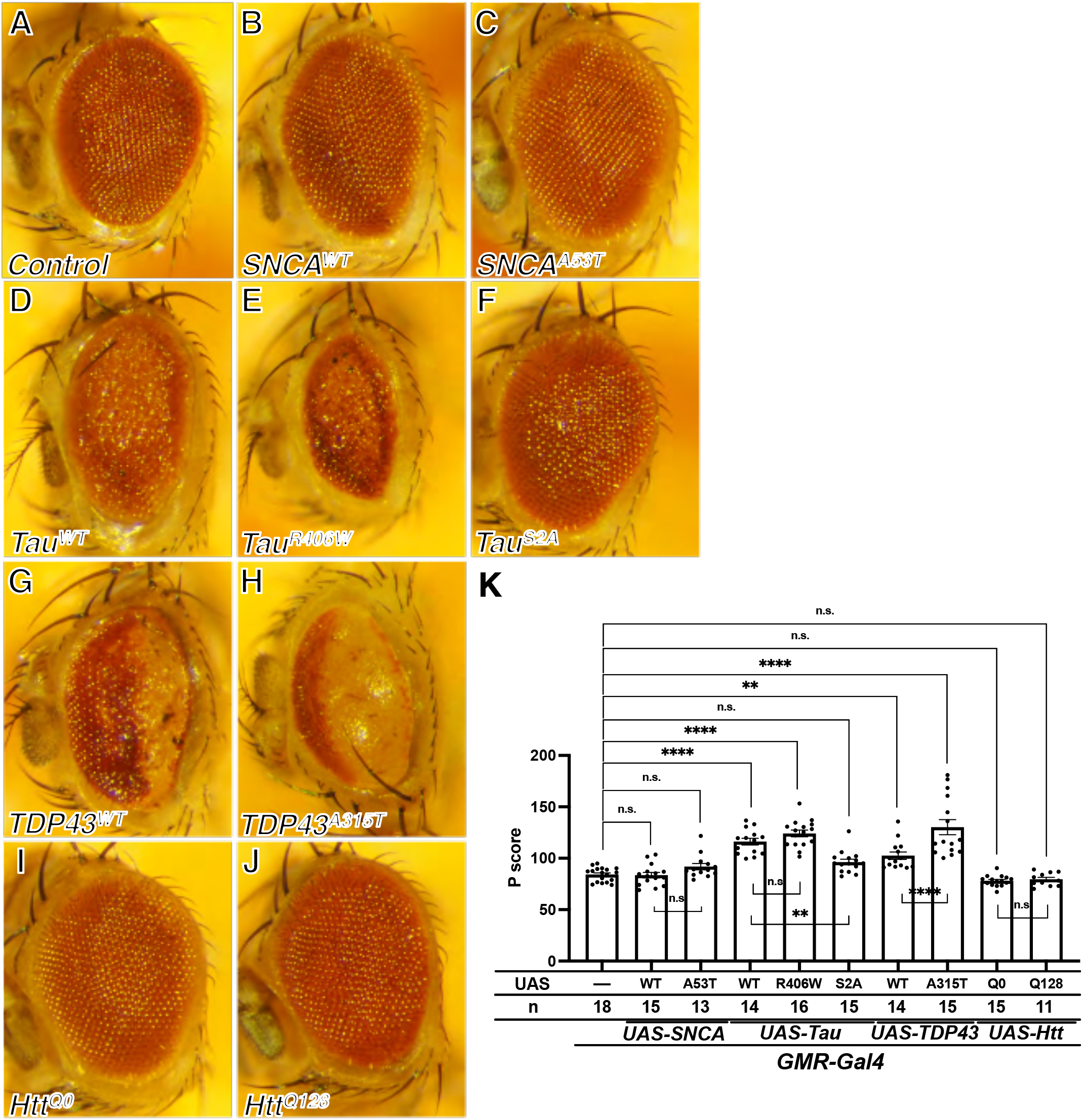
Ectopic expression of causative genes of NDs in fly eye causing REP, but inconsistent with axonal degeneration. (A–J) Representative bright-field microscope images of (A) control or fly eyes overexpressing (B) SNCA^WT^, (C) SNCA ^A53T^, (D) Tau^WT^, (E) Tau^R406W^, (F) Tau^S2A^, (G) TDP-43^WT^, (H) TDP-43^A315T^, (I) HTT^Q0^ and (J) HTT^Q128^ using the GMR-Gal4 driver. (K) Graph representing the phenotypic score (P score) of each genotype calculated using Flynotyper. \*\*\*\**p* < 0.0001, \*\**p* < 0.01 and ns (*p* > 0.05). Data were analysed using multiple comparison ANOVA with Tukey–Kramer *post hoc* tests. Error bars show the standard error of the mean.

## Discussion

We recently developed a method that precisely quantifies the axonal number of retinal R7 neurons by creating a mask which extracts the axon terminals [34]. In the present study, we developed and tested MeDUsA, a software that automatically creates masks and counts axon terminals using a combination of deep learning and Python (Fig. 2). By using this software, the number of R7 axons can be automatically quantified easily and quickly from confocal images. Conventionally, REP is frequently used to assess neurotoxicity in fly studies because it does not require special equipment and can be readily performed; however, we found that the severity of REP is not always consistent with axonal toxicity (Fig. 3 and Fig. 5). Also conditions that induce sporadic progressive axonal degeneration of R7 are not accompanied by cell death [34]. Therefore, we propose that ND research can be conducted more efficiently by combining REP with our method when the focus is axonal degeneration. Although several models for evaluating axonal degeneration in *Drosophila* have been reported [26,27,34], they are technically difficult and exceedingly time-consuming to evaluate axonal degeneration in large-scale experiments such as screening as the evaluation of degeneration in these models is subjective or require manual measurement. However, by using MeDUsA, it is not only possible to identify factors and chemical compounds that inhibit axonal degeneration in the fly model of ND by screening but also to easily evaluate axonal toxicity of new or undiagnosed variants of pathological proteins of ND.

According to our modifier screening using REP to identify genes that suppress TDP-43^G298S^ toxicity, we identified 14 candidate genes. Further investigation focused on axonal degeneration showed that the knockdown of three genes (*mle*, *caz* and *faf*) suppressed toxicity, whereas *Dsk* knockdown promoted it (Fig. 1C, 1D). *Dsk* encodes the cholecystokinin-like neuropeptide Drosulfakinin and has been reported to be involved in feeding behaviour, aggression and larval avoidance [45–48]. A previous study reported that Dsk is also involved in synaptogenesis in the neuromuscular junction in cooperation with a putative cholecystokinin-like receptor [49]. Studies in mouse models have shown that the pathogenic form of TDP-43 has harmful effects on synapses. For example, hyperexcitability has been observed [50] and spine density has been reduced [51]. Therefore, our results suggest that Dsk reduction causes synapse dysfunction and make axons fragile, whereby axonotoxicity of TDP-43^G298S^ is enhanced. Named by its male-specific lethal phenotype in loss-of-function mutants, *mle* (*maleless*) encodes an RNA helicase and is a member of the Male-Specific-Lethal transcription complex, which is involved in dosage compensation in males [52]. The homologue of *mle* in human, *DHX9*, was reported to encode a TDP-43–interacting protein [53]. Furthermore, a previous study reported in a fly model that knockdown of *mle* exacerbated neurodegeneration caused by the expression of expanded UGGAA, which is considered responsible for spinocerebellar ataxia type 31 (SCA31), whereas overexpression of wild-type TDP-43 suppressed expanded UGGAA-induced toxicity [39]. Thus, *mle* may be involved in the RNA-dependent toxicity of TDP-43. *caz* (*cabeza*), which encodes a RNA-binding protein, is a fly homologue of FUS/TLS, which is another major causative gene for ALS. Physical interactions between human FUS/TLS and TDP-43 have been reported in mammalian cultured cells [54, 55], and it has been suggested that FUS/TLS is genetically located downstream of TDP-43 in fly and fish models [56, 57]. Therefore, the attenuation of axonal degeneration by *caz* knockdown may be due to the suppression of excess unknown downstream factors of TDP-43 or *caz* itself or both. The influence of *faf* (fat facets), which encodes a deubiquitylated enzyme and is a fly homologue of *USP9X*, on neurodegeneration varies among fly ND models. Lee et al. reported that FAF enhanced the toxicity of amyloid precursor protein (APP) and Aβ-42. The co-expression of FAF with either APP or Aβ-42 enhanced REP and knockdown of *faf* suppressed the post-synaptic toxicity of APP and Aβ-42 [58]. Another group revealed that reduced levels of *faf* enhanced retinal toxicity of HTT [19]. As FAF deubiquitinates different substrates in these diseases, it may be reflected in the different effects on the toxicity of causative proteins of the ND. Further analysis of these genes would shed light on the molecular mechanisms underlying axonotoxicity by TDP-43^G298S^.

Using the automatic quantification method developed in this study, we evaluated the axonal effects of several representative causative factors of NDs, including αSyn, Tau, TDP-43 and HTT (Fig. 3). Although the pathological significance of these causal factors has been examined in various model organisms, including fly model, few studies have directly visualised and quantitatively evaluated axonal degeneration such as that demonstrated in this study using MeDUsA. Our findings show that MeDUsA is highly extensible and can be used in different NDs to demonstrate how a causative gene affects axonal degeneration.

*SNCA* encodes αSyn, which is highly localised at the pre-synaptic terminal, and is thought to mediate the regulation of synaptic function [59]. Studies using post-mortem brains and primary neurons from patients with PD suggest that αSyn aggregates in axons and causes degeneration, resulting in impaired neuronal function that is propagated to the cell body and leads to neuronal death [60, 61]. The A53T mutation is a well-known toxic mutation of αSyn that aggregates more rapidly and forms fibrils than wild-type, and many studies using animal models and induced pluripotent stem (iPS) cells derived from patients with PD have reported high neurotoxicity [62–64]. In our study, a significant reduction in the number of axons was observed between control and αSyn^A53T^-expressed flies, but no statistically significant difference was observed between wild-type and A53T (Fig. 3K). A possible reason for the non-significant difference is that quantification was performed too early (1-day-old adult), and differences between the wild-type and A53T mutation may be observed if quantified several weeks after eclosion.

Tau is a microtubule-associated protein that binds to microtubules and maintains their stability in neurons. In tauopathies, Tau is thought to be dissociated from microtubules by excessive phosphorylation and aggregate, causing a dying-back pattern of neurodegeneration [65]. Although several studies have reported increased toxicity with the R406W mutation, a change in the phosphorylation status remains controversial. Studies using *in vitro* and patient-derived iPS cells have reported that R406W mutant *tau* is less phosphorylated than wild-type *tau*, whereas excessive phosphorylation has been observed in both murine model and post-mortem patient brain [66–69]. Although previous studies reported that overexpression of Tau^R406W^, which was used in this study, displayed high toxicity in REP compared to Tau^WT^ in *Drosophila* [5, 70], it may be due to positional effects of the UAS insertion site [71]. Consistent with previous reports, we found that Tau^R406W^ overexpression exerted a more toxic effect on axons than Tau^WT^ overexpression (Fig. 3K).

TDP-43 is a highly conserved 43 kDa RNA-binding protein that is a main component of ubiquitinated aggregation in the neurons of patients with ALS and frontotemporal lobar degeneration (FTLD) [72]. Numerous studies that have investigated the physiological function of TDP-43 have revealed that TDP-43 is involved in various aspects of RNA metabolism, and these disturbances may be responsible for the pathogenesis of ALS and FTLD [73]. In many model organisms, both overexpression and loss-of-function of TDP-43 result in reduced longevity and motor function [74–77]. The overexpression of wild-type TDP-43 in *Drosophila* mushroom body neurons of the olfactory memory centre causes axonal degeneration [78], whereas the effects of the A315T mutation on TDP-43 toxicity are controversial in *Drosophila*. Guo et al. reported that TDP-43^A315T^ was more toxic to motor neurons than TDP-43^WT^, whereas Patricia et al. reported that TDP-43^WT^ showed severe toxicity compared to TDP-43^A315T^, except for larval locomotor activity [79, 80]. In our present result of retinal R7 axons, we found that wild-type TDP-43 showed higher axonal toxicity than the A315T mutation (Fig. 3K). Although TDP-43 is primarily expressed in the nucleus, the aggregation of TDP-43 in the cytoplasm causes toxicity. As wild-type TDP-43 may be increased in the cytoplasm if it is overexpressed, it is important to realise that differences in expression levels in each experimental system may contribute to differences in the respective results. TDP-43 may also have different toxic effects on different neuronal cell types.

In Huntington’s disease, CAG repeat expansion of *HTT* produces abnormal RNA and protein, leading to neuronal dysfunction and eventual cell death. The normal allele of *HTT* contains fewer than 26 CAG repeats, whereas 36 repeats or more are associated with Huntington’s disease [81]. A study using a mouse model of HD and human patients indicated that the degeneration of the callosal axon was seen before symptoms were observed, suggesting that the mutant HTT caused dying-back neurodegeneration [82]. Furthermore, CAG repeat number-dependent cytotoxicity has been reported in *Drosophila* [83]. In this study, as in previous reports, HTT^Q128^ showed axonal toxicity but HTT^Q0^ did not (Fig. 3K).

Axonal degeneration is observed not only in NDs but also in Wallerian degeneration (WD), which is the axotomy-induced distal degeneration of an axon, causing a decline in neuronal function. The mechanisms of axonal degeneration in NDs and WD are partly overlapping but not identical. Wlds is a fusion protein that slows WD; it confers a protective effect from degeneration in animal models of progressive motor neuropathy and PD, but does not ameliorate in ALS model [84–86]. These findings suggest that each pathological protein causes axonal degeneration by a different mechanism, but the detailed molecular mechanisms are still poorly understood. However, by using our method, it is anticipated that research focusing on axon degeneration will be facilitated, thereby enabling the elucidation of the pathological mechanism of axonal degeneration by the causative genes of ND and WD. In turn, a better understanding of the underlying molecular mechanisms of axonal degeneration is promising for developing therapies that inhibit or delay the onset of ageing, NDs and WD.

By using deep learning, it was possible to automatically create a mask that extracts axon terminals from a confocal z-stack image. The image processing capabilities afforded by machine learning are powerful, and recently, many quantitative and segmentation methods using machine learning have been published [87–90]. MeDUsA is a Python-based method specifically designed to count axons; however, it only quantifies the presence of axons and does not capture pre-degenerative signs such as swelling or fragmentation of axon terminals. Such changes are currently more precisely captured only with a manual method [34]. An important future step will be to extend MeDUsA to perform a more comprehensive quantitative analysis of morphological and cell biological properties, such as the size and shape of axonal termini, the number or organisation of pre-synapses, of mitochondria or other organelles in axonal terminals, to enable more detailed studies of pathological mechanisms of ND.

## Conclusions

In this study, we developed MeDUsA for automatically quantifying the number of axons in retinal R7 neurons in *Drosophila* with high reliability and robustness. It combines pre-trained deep-learning models with a Python-based quantification system. Using our easy-to-use software, we demonstrated the causative proteins of NDs actually caused the axonal degeneration. We also confirmed that the severity of REP and axonal number were not significantly correlated. MeDUsA is a valuable tool for the unbiased and rapid quantification of axonal degeneration in genetic or pharmacological modifier screening using *Drosophila* as a model.

## Materials and Methods

### Fly Strains

Flies were maintained at 25°C on standard fly food. Female flies were used in all experiments except for those shown in Figure 1 to adjust the number of retinal axons. *40D-UAS* (VDRC ID 60101) was obtained from the Vienna *Drosophila* Resource Center (VDRC) in Vienna, Austria. *GMR-Gal4 (III)* (#8121), *Rh6-Gal4* (#7459), *Brp-FSF-GFP* (#55753), *UAS-myr-RFP* (#7119), *UAS-SNCA* (#8146), *UAS-SNCA^A53T^* (#8148), *UAS-Tau^S2A^* (#51364), *tub-Gal80^TS^* (#7017 and #7019), *UAS-marf RNAi* (#55189), *UAS-opa1 RNAi* (#32358), *GMR-w-RNAi* (#32067), *lexAop-syb-spGFP1-10*, *UAS-CD4-spGFP11* (#64315) and 99 strains for RNAi screening were obtained from the Bloomington *Drosophila* Stock Center (BDSC, Bloomington, IN, USA). Details are provided in Sup. Table 1. *GMR-Gal4(II)* [37, 38] was used for the expression of transgenes in the photoreceptors. *Rh4-LexA* was generated previously [91]. *ortC2b-Gal4* was gifted by Dr. Chi-Hon Lee [92], and *Sens-flippase* [93] was kindly provided by Dr. S. L. Zipursky. *UAS-Tau* [5] and *UAS-Tau^R406W^* [5] were gifted by Dr. M. B. Feany. *UAS-HTT^Q0^* [94]and *UAS-HT*T*^Q128^* [94] were graciously given by the Dr. J. T. Littleton. *UAS-TDP-43(strong)*, *UAS-TDP-43^A315T^*, and *UAS-TDP43^G298S^* were described previously [39, 95].

### Immunohistochemistry and imaging

Immunohistochemistry and sample preparation were performed as described previously [34, 40]. The following antibodies were used: mouse anti-Chaoptin (24B10, 1:25; Developmental Studies Hybridoma Bank, Iowa City, IA, USA), anti-mouse Alexa Fluor 488 (1:400; Thermo Fisher Scientific) and anti-mouse Alexa Fluor 568 (1:400; Thermo Fisher Scientific). The insect pins were used because they are 0.1 mm in diameter and fit the thickness of a fly brain. A coverslip was added and slides were mounted using Vectashield mounting medium (Vector Laboratories). Images were captured using a FV3000 confocal microscope (Olympus, Tokyo, Japan). We then obtained brain images using a confocal microscope set to a 60× immersion objective (1.4 numerical aperture) and a 1× digital zoom. During the scan, we set the step size to 1 μm and generated 60–90 optical sections of the second optic ganglion medulla, including all R7 axon terminals. They are for the images shown in Figure 2, 3 and 6. In Figure 1, samples were scanned using an A1 confocal microscope (Nikon, Tokyo, Japan). Images were processed using either IMARIS 9.6.0 (Bitplane) or Fiji software, which is an open-source image analysis software [96].

### Eye imaging using bright-field microscopy and quantification of morphological eye defects

For light microscopy imaging of adult eyes, 1- to 3-day-old flies, which were reared at 29°C for the expression of αSyn, Tau and HTT and at 25°C for the expression of TDP-43, were immobilised by freezing at −20°C, and separated fly heads were mounted on labelling tape (Shamrock Labels, IL, USA). The flies were then imaged using an OM-D E-M5 digital camera (Olympus) and SZX16 microscope with 6.3× magnification (Olympus). About 20 photographs were taken with the focus shifted slightly from the centre of the eye to the edges. Each slice was depth-synthesised using Photoshop CC 2017 (Adobe). The edges of the eyes were then trimmed and processed using Flynotyper 1.0 to quantitatively assess morphological defects in *Drosophila* eye [24].

### Combined rough eye and axonal degeneration phenotype screening

First, candidate genes for screening were selected as the phenotype in the QuickSearch of FlyBase (https://flybase.org/) and searched for ‘synapse’ in the ‘Tissue/cell affected’ category. The 401 genes hit using this method were further narrowed down using settings of ‘Higher’ and ‘Moderately High’ with ‘Adult Head’ in the modENCODE Expression Data of FlyBase, and 99 genes available for RNAi lines at the BDSC were selected as candidate genes.

For the initial screening, we searched to identify a factor that suppressed REP to serve as an indicator of neurodegeneration. In this study, we used a transgene of TDP-43^G298S^ expressed by the eye-specific *GMR-Gal4* driver to induce REP. We then evaluated each RNAi line of the candidate genes, and the genes in which REP was suppressed were selected. We then imaged the compound eyes of *Drosophila* using an EOS Kiss X4 digital camera (Canon, Tokyo, Japan) and MZ FLIII fluorescence stereo microscope (Leica, Wetzlar, Germany).

Next, as the second screening, R8 axons were visualised to further narrow the candidate factors based on the ability to recover from axonal degeneration. To achieve this, myrRFP was expressed using the *Rh6-Gal4* driver. The genotype was *Sens-flippase/+; Rh6-Gal4/UAS-TDP-43^G298S^; Brp-FSF-GFP, UAS-myr-RFP/UAS-RNAi*. Knockdown was performed on 15 candidate genes that suppressed the REP, and the axons were observed using an A1 confocal microscope (Nikon).

### Training set generation

To quantify both normal and degenerated axons with sufficient accuracy, we used samples with phenotypes of axonal degeneration induced by either light stimulation or mitochondrial dysfunction for machine learning.

*Drosophila* photoreceptors are inherently sensory neurons for light. We found that constant light stimulation caused progressive axonal degeneration [34]. Taking advantage of this phenomenon, we used samples of various time points under constant light from normal states to severe axonal degeneration for machine learning. In detail, experimental samples at 1 (n = 16), 3 (n = 16), 5 (n = 20), 7 (n = 16), 9 (n = 14), 11 (n = 14) and 13 days (n = 13) under constant light were used. Samples (n = 10 and n = 19) were used on day 1 and day 13 in a 12-h light/dark cycle as controls, respectively. The genotype was *GMR-w-RNAi/w-; lexAop-syb-spGFP1-10, UAS-CD4-spGFP11/ Rh4-LexA; ortC2b-Gal4/+*.

For machine learning, we also induced photoreceptor axonal degeneration by knocking down Marf or Opa1, both of which are required for mitochondrial fusion. The genotypes were *GMR-Gal4/40D-UAS; tub-Gal80^ts^/+* (n = 6), *GMR-Gal4/UAS-marf RNAi; tub-Gal80^ts^/+* (n = 9) and *GMR-Gal4/tub-Gal80^ts^; UAS-opa1 RNAi/+* (n = 20). The flies were reared in a permissive temperature (20°C), and after eclosion, knockdown was induced by rearing the flies in a restrictive temperature (29°C) and were dissected 4 weeks later. Samples reared under control conditions at 20°C for 4 weeks after eclosion were also used for machine learning. The genotypes were *GMR-Gal4/40D-UAS; tub-Gal80^ts^/+* (n = 14), *GMR-Gal4/UAS-marf RNAi; tub-Gal80^ts^/+* (n = 20), and *GMR-Gal4/tub-Gal80^ts^; UAS-opa1 RNAi/+* (n = 10).

All samples were scanned using a confocal microscope. We then manually created a mask covering the axonal terminal and used the scanning data and the mask as a training set.

### Surface mask prediction and axon terminal detection

A variant of 2D-U-Net was used for surface mask prediction. The original images were N × 512 × 512 in size for each sample, and for each z-slice, the image before and after it (z − 1, z and z + 1) were combined to form the N × 512 × 512 × 3 size. As the terminal slice does not have z − 1 or z + 1, a blank image was used instead. For training, we used mask images of size N × 512 × 512 × 3, and for inference, we obtained mask prediction images of size N × 512 × 512 by discarding the channels corresponding to each z − 1 and z + 1 from the N × 512 × 512 × 3 output and retaining only z. As post-processing, opening and closing operations were used to exclude small blobs and to fill in holes. Then, from the original image, only the area corresponding to the obtained surface mask image was extracted. To detect the axon terminal, the background image was first created by morphological reconstruction and then filtered regional maxima by subtracting it from the original image to remove the background. Next, the image was binarised by adaptive thresholding, and 3D watershed was performed using the peak obtained by calculating the Euclidean distance as a seed. Finally, blobs below 20 voxels were excluded. These processes were calculated using Python (v3.6.7), NumPy (v1.17.3), TensorFlow (v1.13.2), scikit-image (v0.16.2), SciPy (v1.3.2) and OpenCV (v4.2.0).

### Experimental design and statistical analyses

Experimental analyses were performed using Prism 8 (GraphPad Software, San Diego, CA, USA). All quantifications were performed by experimenters who were blind to the genotype. Data were analysed using multiple comparison ANOVA with Tukey–Kramer post hoc tests or unpaired t-test with Mann–Whitney test, as noted in the Results section. The null hypothesis was rejected at a 0.05 level of significance.

## List of Abbreviations

MeDUsA: a method for quantification of degeneration using fly axons
ND: Neurodegenerative disease
REP: rough eye phenotype
SCA3: Spinocerebellar ataxia type 3
PD: Parkinson’s disease
ALS: amyotrophic lateral sclerosis
CNN: convolutional neural network
RNAi: RNA interference
SCA31: spinocerebellar ataxia type 31
FTLD: frontotemporal lobar degeneration
WD: Wallerian degeneration

## Declarations

Ethics approval and consent to participate Not applicable

## Consent for publication

Not applicable

## Availability of data and materials

The software documentation for MeDUsA can be found at https://github.com/SugieLab/MeDUsA.

The datasets used and/or analyzed during the current study are available from the corresponding authors on reasonable request.

## Authors’ contributions

YNI and AS designed and organized the study.

YNI and AS performed immunohistochemistry, and eye imaging.

YNI, AS and KD performed data analysis.

HK established the software.

JO, SHS, and TS designed the screen experiment and JO performed the experiment and data analysis.

YNA generated transgenic flies.

YNI, HK, KD, GT and AS wrote the manuscript

All authors read and approved the final manuscript.

## Acknowledgements

We would like to acknowledge that Dr. Zipursky, Dr. Feany, Dr. Lee and Dr. Littleton have provided us with transgenic fly strains. We would like to thank Ms. Nozaki for helping us to quantify REP.

## Reference

1. Dugger BN, Dickson DW. Pathology of Neurodegenerative Diseases. Csh Perspect Biol. 2017;9:a028035.

2. Warrick JM, Paulson HL, Gray-Board GL, Bui QT, Fischbeck KH, Pittman RN, et al. Expanded Polyglutamine Protein Forms Nuclear Inclusions and Causes Neural Degeneration in Drosophila. Cell. 1998;93:939–49.

3. Jackson GR, Salecker I, Dong X, Yao X, Arnheim N, Faber PW, et al. Polyglutamine-Expanded Human Huntingtin Transgenes Induce Degeneration of Drosophila Photoreceptor Neurons. Neuron. 1998;21:633–42.

4. Feany MB, Bender WW. A Drosophila model of Parkinson’s disease. Nature. 2000;404:394–8.

5. Wittmann CW, Wszolek MF, Shulman JM, Salvaterra PM, Lewis J, Hutton M, et al. Tauopathy in Drosophila: neurodegeneration without neurofibrillary tangles. Science. 2001;293:711–4.

6. Chouhan AK, Guo C, Hsieh Y-C, Ye H, Senturk M, Zuo Z, et al. Uncoupling neuronal death and dysfunction in Drosophila models of neurodegenerative disease. Acta Neuropathologica Commun. 2016;4:62.

7. Elden AC, Kim H-J, Hart MP, Chen-Plotkin AS, Johnson BS, Fang X, et al. Ataxin-2 intermediate-length polyglutamine expansions are associated with increased risk for ALS. Nature. 2010;466:1069–75.

8. Brand AH, Perrimon N. Targeted gene expression as a means of altering cell fates and generating dominant phenotypes. Development. 1993;118:401–15.

9. Blard O, Feuillette S, Bou J, Chaumette B, Frébourg T, Campion D, et al. Cytoskeleton proteins are modulators of mutant tau-induced neurodegeneration in Drosophila. Hum Mol Genet. 2007;16:555–66.

10. Chen X, Li Y, Huang J, Cao D, Yang G, Liu W, et al. Study of tauopathies by comparing Drosophila and human tau in Drosophila. Cell Tissue Res. 2007;329:169–78.

11. Ambegaokar SS, Jackson GR. Functional genomic screen and network analysis reveal novel modifiers of tauopathy dissociated from tau phosphorylation. Hum Mol Genet. 2011;20:4947–77.

12. M’Angale PG, Staveley BE. The Bcl-2 homologue Buffy rescues α-synuclein-induced Parkinson disease-like phenotypes in Drosophila. Bmc Neurosci. 2016;17:24.

13. Alexopoulou Z, Lang J, Perrett RM, Elschami M, Hurry MED, Kim HT, et al. Deubiquitinase Usp8 regulates α-synuclein clearance and modifies its toxicity in Lewy body disease. Proc National Acad Sci. 2016;113:E4688–97.

14. Davies SE, Hallett PJ, Moens T, Smith G, Mangano E, Kim HT, et al. Enhanced ubiquitin-dependent degradation by Nedd4 protects against α-synuclein accumulation and toxicity in animal models of Parkinson’s disease. Neurobiol Dis. 2014;64:79–87.

15. Miura E, Hasegawa T, Konno M, Suzuki M, Sugeno N, Fujikake N, et al. VPS35 dysfunction impairs lysosomal degradation of α-synuclein and exacerbates neurotoxicity in a Drosophila model of Parkinson’s disease. Neurobiol Dis. 2014;71:1–13.

16. Zhan L, Hanson KA, Kim SH, Tare A, Tibbetts RS. Identification of Genetic Modifiers of TDP-43 Neurotoxicity in Drosophila. Plos One. 2013;8:e57214.

17. Kim H-J, Raphael AR, LaDow ES, McGurk L, Weber RA, Trojanowski JQ, et al. Therapeutic modulation of eIF2α phosphorylation rescues TDP-43 toxicity in amyotrophic lateral sclerosis disease models. Nat Genet. 2014;46:152–60.

18. Calpena E, Amo VL del, Chakraborty M, Llamusí B, Artero R, Espinós C, et al. The Drosophila junctophilin gene is functionally equivalent to its four mammalian counterparts and is a modifier of a Huntingtin poly-Q expansion and the Notch pathway. Dis Model Mech. 2017;11:dmm029082.

19. Kaltenbach LS, Orr H, Romero E, Becklin RR, Chettier R, Bell R, et al. Huntingtin interacting proteins are genetic modifiers of neurodegeneration. PLoS Genetics. 2007;3:e82.

20. Doumanis J, Wada K, Kino Y, Moore AW, Nukina N. RNAi Screening in Drosophila Cells Identifies New Modifiers of Mutant Huntingtin Aggregation. Plos One. 2009;4:e7275.

21. Shim K-H, Kim S-H, Hur J, Kim D-H, Demirev AV, Yoon S-Y. Small-molecule drug screening identifies drug Ro 31-8220 that reduces toxic phosphorylated tau in Drosophila melanogaster. Neurobiol Dis. 2019;130:104519.

22. Lin Y, Xue J, Deng J, He H, Luo S, Chen J, et al. Neddylation activity modulates the neurodegeneration associated with fragile X associated tremor/ataxia syndrome (FXTAS) through regulating Sima. Neurobiol Dis. 2020;143:105013.

23. Diez-Hermano S, Valero J, Rueda C, Ganfornina MD, Sanchez D. An automated image analysis method to measure regularity in biological patterns: a case study in a Drosophila neurodegenerative model. Molecular neurodegeneration. 2015;10:9.

24. Iyer J, Wang Q, Le T, Pizzo L, Grönke S, Ambegaokar SS, et al. Quantitative Assessment of Eye Phenotypes for Functional Genetic Studies Using Drosophila melanogaster. G3 (Bethesda, Md). 2016;6:1427–37.

25. McGurk L, Berson A, Bonini NM. Drosophila as an In Vivo Model for Human Neurodegenerative Disease. Genetics. 2015;201:377–402.

26. Bhattacharya MRC, Gerdts J, Naylor SA, Royse EX, Ebstein SY, Sasaki Y, et al. A model of toxic neuropathy in Drosophila reveals a role for MORN4 in promoting axonal degeneration. Journal of Neuroscience. 2012;32:5054–61.

27. Sreedharan J, Neukomm LJ, Brown RH, Freeman MR. Age-Dependent TDP-43-Mediated Motor Neuron Degeneration Requires GSK3, hat-trick, and xmas-2. Curr Biol. 2015;25:2130–6.

28. Moen E, Bannon D, Kudo T, Graf W, Covert M, Valen DV. Deep learning for cellular image analysis. Nat Methods. 2019;16:1233–46.

29. Ronneberger O, Fischer P, Brox T. U-Net: Convolutional Networks for Biomedical Image Segmentation. Lect Notes Comput Sc. 2015;234–41.

30. Çiçek Ö, Abdulkadir A, Lienkamp SS, Brox T, Ronneberger O. 3D U-Net: Learning Dense Volumetric Segmentation from Sparse Annotation. Medical Image Computing and Computer-Assisted Intervention – MICCAI 2016, 19th International Conference, Athens, Greece, October 17-21, 2016, Proceedings, Part II. 2016. p. 424–32.

31. Falk T, Mai D, Bensch R, Çiçek Ö, Abdulkadir A, Marrakchi Y, et al. U-Net: deep learning for cell counting, detection, and morphometry. Nat Methods. 2019;16:67–70.

32. Fischer CA, Besora-Casals L, Rolland SG, Haeussler S, Singh K, Duchen M, et al. MitoSegNet: Easy-to-use Deep Learning Segmentation for Analyzing Mitochondrial Morphology. iScience. 2020;23:101601.

33. Long F. Microscopy cell nuclei segmentation with enhanced U-Net. Bmc Bioinformatics. 2020;21:8.

34. Richard M, Doubková K, Nitta Y, Kawai H, Sugie A, Tavosanis G. A quantitative model of sporadic axonal degeneration in the Drosophila visual system. bioRxiv. 2021;

35. Sang T-K, Jackson GR. Drosophila models of neurodegenerative disease. Neurorx. 2005;2:438–46.

36. Deerlin VMV, Leverenz JB, Bekris LM, Bird TD, Yuan W, Elman LB, et al. TARDBP mutations in amyotrophic lateral sclerosis with TDP-43 neuropathology: a genetic and histopathological analysis. Lancet Neurology. 2008;7:409–16.

37. Newsome TP, Asling B, Dickson BJ. Analysis of Drosophila photoreceptor axon guidance in eye-specific mosaics. Development. 2000;127:851–60.

38. Newsome TP, Schmidt S, Dietzl G, Keleman K, Asling B, Debant A, et al. Trio combines with dock to regulate Pak activity during photoreceptor axon pathfinding in Drosophila. Cell. 2000;101:283–94.

39. Ishiguro T, Sato N, Ueyama M, Fujikake N, Sellier C, Kanegami A, et al. Regulatory Role of RNA Chaperone TDP-43 for RNA Misfolding and Repeat-Associated Translation in SCA31. Neuron. 2017;94:108–124.e7.

40. Sugie A, Möhl C, Hakeda-Suzuki S, Matsui H, Suzuki T, Tavosanis G. Analyzing Synaptic Modulation of Drosophila melanogaster Photoreceptors after Exposure to Prolonged Light. J Vis Exp. 2017;

41. Spillantini MG, Schmidt ML, Lee VM-Y, Trojanowski JQ, Jakes R, Goedert M. α-Synuclein in Lewy bodies. Nature. 1997;388:839–40.

42. Polymeropoulos MH, Lavedan C, Leroy E, Ide SE, Dehejia A, Dutra A, et al. Mutation in the α-Synuclein Gene Identified in Families with Parkinson9s Disease. Science. 1997;276:2045–7.

43. Hutton M, Lendon CL, Rizzu P, Baker M, Froelich S, Houlden H, et al. Association of missense and 5′-splice-site mutations in tau with the inherited dementia FTDP-17. Nature. 1998;393:702–5.

44. Gitcho MA, Baloh RH, Chakraverty S, Mayo K, Norton JB, Levitch D, et al. TDP- 43 A315T mutation in familial motor neuron disease. Ann Neurol. 2008;63:535–8.

45. Chen X, Peterson J, Nachman RJ, Ganetzky B. Drosulfakinin activates CCKLR-17D1 and promotes larval locomotion and escape response in Drosophila. Fly. 2012;6:290–7.

46. Williams MJ, Goergen P, Rajendran J, Zheleznyakova G, Hägglund MG, Perland E, et al. Obesity-Linked Homologues TfAP-2 and Twz Establish Meal Frequency in Drosophila melanogaster. Plos Genet. 2014;10:e1004499.

47. Agrawal P, Kao D, Chung P, Looger LL. The neuropeptide Drosulfakinin regulates social isolation-induced aggression in Drosophila. J Exp Biol. 2020;223:jeb207407.

48. Wu F, Deng B, Xiao N, Wang T, Li Y, Wang R, et al. A neuropeptide regulates fighting behavior in Drosophila melanogaster. Elife. 2020;9:e54229.

49. Chen X, Ganetzky B. A neuropeptide signaling pathway regulates synaptic growth in Drosophila. J Cell Biol. 2012;196:529–43.

50. Fogarty MJ, Klenowski PM, Lee JD, Drieberg-Thompson JR, Bartlett SE, Ngo ST, et al. Cortical synaptic and dendritic spine abnormalities in a presymptomatic TDP-43 model of amyotrophic lateral sclerosis. Sci Rep-uk. 2016;6:37968.

51. Jiang T, Handley E, Brizuela M, Dawkins E, Lewis KEA, Clark RM, et al. Amyotrophic lateral sclerosis mutant TDP-43 may cause synaptic dysfunction through altered dendritic spine function. Dis Model Mech. 2019;12:dmm038109.

52. Breen TR, Lucchesi JC. Analysis of the dosage compensation of a specific transcript in Drosophila melanogaster. Genetics. 1986;112:483–91.

53. Freibaum BD, Chitta RK, High AA, Taylor JP. Global Analysis of TDP-43 Interacting Proteins Reveals Strong Association with RNA Splicing and Translation Machinery. J Proteome Res. 2010;9:1104–20.

54. Kim SH, Shanware NP, Bowler MJ, Tibbetts RS. Amyotrophic Lateral Sclerosis-associated Proteins TDP-43 and FUS/TLS Function in a Common Biochemical Complex to Co-regulate HDAC6 mRNA*. J Biol Chem. 2010;285:34097–105.

55. Ling S-C, Albuquerque CP, Han JS, Lagier-Tourenne C, Tokunaga S, Zhou H, et al. ALS-associated mutations in TDP-43 increase its stability and promote TDP-43 complexes with FUS/TLS. Proc National Acad Sci. 2010;107:13318–23.

56. Wang J-W, Brent JR, Tomlinson A, Shneider NA, McCabe BD. The ALS-associated proteins FUS and TDP-43 function together to affect Drosophila locomotion and life span. J Clin Invest. 2011;121:4118–26.

57. Kabashi E, Bercier V, Lissouba A, Liao M, Brustein E, Rouleau GA, et al. FUS and TARDBP but Not SOD1 Interact in Genetic Models of Amyotrophic Lateral Sclerosis. Plos Genet. 2011;7:e1002214.

58. Lee S, Wang J-W, Yu W, Lu B. Phospho-dependent ubiquitination and degradation of PAR-1 regulates synaptic morphology and tau-mediated Aβ toxicity in Drosophila. Nature Communications. 2012;3:1312–12.

59. Burré J, Sharma M, Südhof TC. Cell Biology and Pathophysiology of α-Synuclein. Csh Perspect Med. 2018;8:a024091.

60. Orimo S, Uchihara T, Nakamura A, Mori F, Kakita A, Wakabayashi K, et al. Axonal α-synuclein aggregates herald centripetal degeneration of cardiac sympathetic nerve in Parkinson’s disease. Brain. 2008;131:642–50.

61. Volpicelli-Daley LA, Luk KC, Patel TP, Tanik SA, Riddle DM, Stieber A, et al. Exogenous α-Synuclein Fibrils Induce Lewy Body Pathology Leading to Synaptic Dysfunction and Neuron Death. Neuron. 2011;72:57–71.

62. Lashuel HA, Overk CR, Oueslati A, Masliah E. The many faces of α-synuclein: from structure and toxicity to therapeutic target. Nature Publishing Group. 2013;14:38–48.

63. Kouroupi G, Taoufik E, Vlachos IS, Tsioras K, Antoniou N, Papastefanaki F, et al. Defective synaptic connectivity and axonal neuropathology in a human iPSC-based model of familial Parkinson’s disease. Proc National Acad Sci. 2017;114:E3679–88.

64. Bengoa-Vergniory N, Roberts RF, Wade-Martins R, Alegre-Abarrategui J. Alpha-synuclein oligomers: a new hope. Acta Neuropathologica. 2017;134:819–38.

65. Kolarova M, García-Sierra F, Bartos A, Ricny J, Ripova D. Structure and Pathology of Tau Protein in Alzheimer Disease. Int J Alzheimer’s Dis. 2012;2012:731526.

66. Dayanandan R, Slegtenhorst MV, Mack TGA, Ko L, Yen S-H, Leroy K, et al. Mutations in tau reduce its microtubule binding properties in intact cells and affect its phosphorylation. Febs Lett. 1999;446:228–32.

67. Miyasaka T, Morishima-Kawashima M, Ravid R, Heutink P, Swieten JC van, Nagashima K, et al. Molecular Analysis of Mutant and Wild-Type Tau Deposited in the Brain Affected by the FTDP-17 R406W Mutation. Am J Pathology. 2001;158:373–9.

68. Ikeda M, Kawarai T, Kawarabayashi T, Matsubara E, Murakami T, Sasaki A, et al. Accumulation of Filamentous Tau in the Cerebral Cortex of Human Tau R406W Transgenic Mice. Am J Pathology. 2005;166:521–31.

69. Nakamura M, Shiozawa S, Tsuboi D, Amano M, Watanabe H, Maeda S, et al. Pathological Progression Induced by the Frontotemporal Dementia-Associated R406W Tau Mutation in Patient-Derived iPSCs. Stem Cell Rep. 2019;13:684–99.

70. Jackson GR, Wiedau-Pazos M, Sang T-K, Wagle N, Brown CA, Massachi S, et al. Human Wild-Type Tau Interacts with wingless Pathway Components and Produces Neurofibrillary Pathology in Drosophila. Neuron. 2002;34:509–19.

71. Povellato G, Tuxworth RI, Hanger DP, Tear G. Modification of the Drosophila model of in vivo Tau toxicity reveals protective phosphorylation by GSK3β. Biol Open. 2013;3:1–11.

72. Neumann M, Sampathu DM, Kwong LK, Truax AC, Micsenyi MC, Chou TT, et al. Ubiquitinated TDP-43 in Frontotemporal Lobar Degeneration and Amyotrophic Lateral Sclerosis. Science. 2006;314:130–3.

73. Prasad A, Bharathi V, Sivalingam V, Girdhar A, Patel BK. Molecular Mechanisms of TDP-43 Misfolding and Pathology in Amyotrophic Lateral Sclerosis. Front Mol Neurosci. 2019;12:25.

74. Kabashi E, Lin L, Tradewell ML, Dion PA, Bercier V, Bourgouin P, et al. Gain and loss of function of ALS-related mutations of TARDBP (TDP-43) cause motor deficits in vivo. Hum Mol Genet. 2010;19:671–83.

75. Xu Y-F, Gendron TF, Zhang Y-J, Lin W-L, D’Alton S, Sheng H, et al. Wild-Type Human TDP-43 Expression Causes TDP-43 Phosphorylation, Mitochondrial Aggregation, Motor Deficits, and Early Mortality in Transgenic Mice. J Neurosci. 2010;30:10851–9.

76. Ash PEA, Zhang Y-J, Roberts CM, Saldi T, Hutter H, Buratti E, et al. Neurotoxic effects of TDP-43 overexpression in C. elegans. Hum Mol Genet. 2010;19:3206–18.

77. Diaper DC, Adachi Y, Sutcliffe B, Humphrey DM, Elliott CJH, Stepto A, et al. Loss and gain of Drosophila TDP-43 impair synaptic efficacy and motor control leading to age-related neurodegeneration by loss-of-function phenotypes. Hum Mol Genet. 2013;22:1539–57.

78. Li Y, Ray P, Rao EJ, Shi C, Guo W, Chen X, et al. A Drosophila model for TDP-43 proteinopathy. Proc National Acad Sci. 2010;107:3169–74.

79. Guo W, Chen Y, Zhou X, Kar A, Ray P, Chen X, et al. An ALS-associated mutation affecting TDP-43 enhances protein aggregation, fibril formation and neurotoxicity. Nat Struct Mol Biol. 2011;18:822–30.

80. Estes PS, Boehringer A, Zwick R, Tang JE, Grigsby B, Zarnescu DC. Wild-type and A315T mutant TDP-43 exert differential neurotoxicity in a Drosophila model of ALS. Hum Mol Genet. 2011;20:2308–21.

81. Roos RA. Huntington’s disease: a clinical review. Orphanet J Rare Dis. 2010;5:40.

82. Gatto RG, Chu Y, Ye AQ, Price SD, Tavassoli E, Buenaventura A, et al. Analysis of YFP(J16)-R6/2 reporter mice and postmortem brains reveals early pathology and increased vulnerability of callosal axons in Huntington’s disease. Hum Mol Genet. 2015;24:5285–98.

83. Zhang S, Binari R, Zhou R, Perrimon N. A Genomewide RNA Interference Screen for Modifiers of Aggregates Formation by Mutant Huntingtin in Drosophila. Genetics. 2010;184:1165–79.

84. Ferri A, Sanes JR, Coleman MP, Cunningham JM, Kato AC. Inhibiting Axon Degeneration and Synapse Loss Attenuates Apoptosis and Disease Progression in a Mouse Model of Motoneuron Disease. Curr Biol. 2003;13:669–73.

85. Sajadi A, Schneider BL, Aebischer P. Wlds-Mediated Protection of Dopaminergic Fibers in an Animal Model of Parkinson Disease. Curr Biol. 2004;14:326–30.

86. Velde CV, Garcia ML, Yin X, Trapp BD, Cleveland DW. The neuroprotective factor Wlds does not attenuate mutant SOD1-mediated motor neuron disease. Neuromol Med. 2004;5:193–203.

87. Haberl MG, Churas C, Tindall L, Boassa D, Phan S, Bushong EA, et al. CDeep3M—Plug-and-Play cloud-based deep learning for image segmentation. Nat Methods. 2018;15:677–80.

88. Diez-Hermano S, Ganfornina MD, Vegas-Lozano E, Sanchez D. Machine Learning Representation of Loss of Eye Regularity in a Drosophila Neurodegenerative Model. Front Neurosci-switz. 2020;14:516.

89. Kayasandik CB, Ru W, Labate D. A multistep deep learning framework for the automated detection and segmentation of astrocytes in fluorescent images of brain tissue. Sci Rep-uk. 2020;10:5137.

90. Fogo GM, Anzell AR, Maheras KJ, Raghunayakula S, Wider JM, Emaus KJ, et al. Machine learning-based classification of mitochondrial morphology in primary neurons and brain. Sci Rep-uk. 2021;11:5133.

91. Berger-Müller S, Sugie A, Takahashi F, Tavosanis G, Hakeda-Suzuki S, Suzuki T. Assessing the Role of Cell-Surface Molecules in Central Synaptogenesis in the Drosophila Visual System. Plos One. 2013;8:e83732.

92. Gao S, Takemura S, Ting C-Y, Huang S, Lu Z, Luan H, et al. The Neural Substrate of Spectral Preference in Drosophila. Neuron. 2008;60:328–42.

93. Chen Y, Akin O, Nern A, Tsui CYK, Pecot MY, Zipursky SL. Cell-type-Specific Labeling of Synapses In Vivo through Synaptic Tagging with Recombination. Neuron. 2014;81:280–93.

94. Lee W-CM, Yoshihara M, Littleton JT. Cytoplasmic aggregates trap polyglutamine-containing proteins and block axonal transport in a Drosophila model of Huntington’s disease. P Natl Acad Sci Usa. 2004;101:3224–9.

95. Saitoh Y, Fujikake N, Okamoto Y, Popiel HA, Hatanaka Y, Ueyama M, et al. p62 Plays a Protective Role in the Autophagic Degradation of Polyglutamine Protein Oligomers in Polyglutamine Disease Model Flies*. J Biol Chem. 2015;290:1442–53.

96. Schindelin J, Arganda-Carreras I, Frise E, Kaynig V, Longair M, Pietzsch T, et al. Fiji: an open-source platform for biological-image analysis. Nature Methods. 2012;9:676–82.

